# Distinctive neurophysiological correlates of sound onset and offset perception in humans

**DOI:** 10.1101/2025.07.23.666338

**Authors:** Fatima Ali, Gabriela Bury, Adèle Simon, Jennifer F. Linden

## Abstract

Accurate detection of sound onsets and offsets is vital for speech perception. Sound-onset event-related potentials (ERPs) have been well-characterised in electroencephalography (EEG) studies, but the characteristics of sound-offset ERPs have often been obscured by temporal confounds in experimental design. Here, two EEG studies were conducted in human listeners performing an interval duration discrimination task with (i) noise intervals in silence or (ii) silent intervals (gaps) in noise. Stimuli used in the active task were also presented under passive listening conditions. This design enabled us to investigate whether features of the offset ERP could be masked by task demands, and whether the usual temporal precedence of the onset cue in a sound influences the relative magnitude and shape of onset and offset ERPs. The morphology of the offset ERP was distinct from that of the onset ERP in both noise and silent interval duration discrimination tasks, even though the roles of onset and offset cues as initial versus final markers of interval duration were reversed. This observation corroborates evidence from animal studies that there are fundamental differences in brain mechanisms of onset and offset perception. In all experimental conditions, the amplitude of the offset ERP was one-third to one-half that of the onset ERP. Differences between active and passive listening conditions were largely explained by enhancement of ERPs for whichever cue (onset or offset) marked the end of the intervals compared in the duration discrimination task. However, this context dependence emerged in earlier ERP waves for offset than onset responses.

**Key points summary:**

- Onset and offset event-related potentials (ERPs) differ in morphology and amplitude
- For both onset and offset ERPs, P200 and later waves depend on behavioural context
- For offset ERPs only, P100/N100 waves also depend on behavioural context
- Engagement in duration discrimination alters ERPs primarily for end-interval cues

## Introduction

Accurate detection of fast temporal fluctuations in sound is critical for perception of speech, especially in varying background noise. The transients of these temporal fluctuations – sound onsets and offsets – mark the boundaries between silence and sound intervals. Neurobiological studies in animals have recently elucidated distinct responses, neural mechanisms and pathways for central auditory processing of sound offsets and onsets (Scholl et al., 2010; Anderson & Linden, 2016; Kopp-Scheinpflug et al., 2018). However, the implications of this functional dissociation for human perception are not known. Most auditory perceptual and neurophysiological studies in humans have focused exclusively on sound onsets as the vital temporal cue, or used short stimuli that preclude separation of brain responses to sound onsets and offsets.

Brain-behaviour relationships in speech perception cannot be fully understood without consideration of the offset cue. Onset and offset cues occur in similar proportions in naturalistic speech sounds, and both cues are critical for discriminating durations of intervals of sound and silence that allow us to distinguish between particular pairs of speech sounds (Dorman et al., 1979; Allen & Li, 2009). Moreover, deficits in auditory temporal processing can arise from specific impairments in sound-offset sensitivity alone in animal models (Anderson & Linden, 2016). In humans, impaired auditory temporal processing has been linked to difficulties in understanding speech in noisy environments, even in those with normal hearing thresholds (American Academy of Audiology [AAA], 2010; Eggermont, 2015, Hind et al., 2011; Iliadou et al., 2017). It is possible that specific impairments in sound-offset sensitivity could contribute to auditory temporal processing difficulties in humans that are currently difficult to diagnose and poorly understood.

EEG is an ideal technology to use for investigating the role of offsets and onsets in human auditory perception because it can be used to detect brain responses to stimulus events with very high temporal resolution; moreover, EEG technology is non-invasive, relatively inexpensive, and already widely available in audiology clinics. Auditory event-related potentials are typically largest in the fronto-central regions of the cortex due to dispersion of EEG signals by the head tissues, although the N100 and P200 waves have neural generators in the auditory cortex (Ross et al., 2009). Early studies using tone bursts showed that the duration of tone stimuli required to elicit a distinct offset ERP was in the range 0.8 to 1.5s, and the offset ERP evoked by long tones was similar in N100/P200 morphology to the onset response (Rose and Malone, 1965; Davis and Zerlin, 1966; Onishi and Davis, 1968). With equal preceding intervals of sound or silence, the offset ERP amplitude was no more than half of the onset ERP amplitude (Onishi and Davis, 1968; Spychala et al., 1969); however, offset responses could be made more comparable in size to onset responses by using long tones compared to short gaps to increase the novelty of the end of the tone relative to its beginning (Pfefferbaum et al., 1971). Although most studies showed no latency difference between onset and offset ERPs, two studies reported an earlier N100 peak for offsets compared to onsets, most evident at high intensities (Onishi and Davis, 1968; Johannsen et al., 1972). Moreover, magnetoencephalography (MEG) methods with greater spatial resolution than EEG have shown that though offset and onset ERPs have many similarities including scalp distributions, there is no P50 for the offset waveform, suggesting that P50 cannot be causal to the N100 component in the offset ERP (Hari et al., 1987; Pantev et al., 1996; Yamashiro et al., 2009; Pratt et al., 2008; Zhang et al., 2016).

Onsets have an additional confound in the novelty of the physical input whereas offsets are a temporal cue for the end of a particular sound (Herdener et al., 2007). Effects of attention on the auditory ERP might therefore be expected to be more pronounced for sound onsets than offsets because of their greater stimulus novelty. Hillyard et al. showed that the N100 component of the onset ERP was enhanced by selective attention to the auditory stimulus (Hillyard et al., 1973). In a group of studies by Pratt et al, no such effect of attention was found on early waves of the offset ERP, only on the much later P300 wave (Pratt et al., 2005; Pratt et al., 2007). A more recent study reported delayed N100 latencies for offset ERPs when the difficulty of a tone duration discrimination task was increased, suggesting that offset responses might be slowed by increased attentional load (Volosin & Horvath, 2020). However, in this and other studies of ERPs measured during sound duration discrimination tasks, the offset cue was always closer in time to a behavioural decision about duration than the onset cue; therefore, general effects of attention on brain responses to stimulus transients could not be distinguished from more temporally specific effects of engagement in a duration discrimination task.

Here we revisited the question of whether sound offsets and onsets evoke different ERP waveforms with different dependence on attention and/or behavioural task. We sought to tackle four common limitations of previous EEG studies of sound offsets. First, some previous studies have used stimulus durations too short to allow clean separation of offset and onset ERP profiles, instead relying on difference waveforms to extract the response to the offset alone or interpreting the overlap as an interaction between onset and offset (Hillyard and Picton, 1978). We used stimuli of sufficient duration to elicit both onset and offset responses without overlap. Second, many studies have not attempted to control participant attention with a behavioural task; it is possible in those studies that the participants were not attending to the onsets or offsets at all, or attending to them differently. In the current study, duration discrimination tasks requiring precise attention to both onset and offset cues were used. Third, some previous studies evoked onset and offset responses using stimuli with very few possible durations, increasing the temporal predictability of the transient events. By using a large range of randomised durations for the duration discrimination task, we minimised contingent negative variation related to anticipation prior to the event (Walter et al., 1964; Kononowicz & Penney, 2016) and reduced the likelihood of a sustained anticipatory response that could overlap with the transient response (Picton et al., 1978). Last, in some studies, the onset cue was always prior to the offset cue, a temporal confound that could influence the relative salience of the two events as well as their proximity to duration decisions. By reversing the progression of these cues so that the offset has the functional role of indicating an initial change from the background perception and the start of the duration interval, we were able to assess the contribution of cue order to onset and offset ERPs.

We measured onset and offset ERPs while participants performed one of two long-duration discrimination tasks or listened passively to the same stimuli. One task involved discrimination of the durations of long noises presented on a silent background; the other required discrimination of the durations of long silent gaps presented on a noise background. We used white noises instead of tones to minimise spectral splatter confounds and to maximise the potential clinical utility of the stimulus paradigm. Both tasks required specific attention and precision in detecting onsets and offsets because durations were randomised over a continuous rather than discretized range. By reversing the order of the onset/offset cues to test the figure-ground relationship of the responses, and by measuring onset and offset ERPs in both active duration discrimination and passive listening conditions in each participant, we identified distinctive and reliable features of the offset-related ERP profile and resolved open questions about the task-dependence of offset ERPs compared to onset ERPs.

## Methods

Experimental procedures were approved by the UCL Research Ethics Committee (10433/001). All participants in this study were aged between 18 and 44 years, healthy, and had no history of developmental, cognitive, or hearing problems. All reported that they had not been exposed to any prolonged loud noise in the previous 24 hours. They had normal hearing according to standard pure tone audiometry measures tested at the start of the experiment. All subjects were paid for their participation.

Two studies were conducted on two largely distinct groups of participants (Study 1: n=12: 8 f, 4 m, age range 18-44, mean 28.8 ± 8.1 y; Study 2: n=11: 6 f, 5 m, age range 19-39, mean 25.9 ± 6.0 y). Only three participants took part in both Study 1 and 2. In Study 1, the participants were required to discriminate between two noises in background silence, while in Study 2, the participants were required to discriminate between two gaps in background noise (see Figure 1). This design enabled us to investigate the role of onsets and offsets with reversed functions of signalling the start or end of the interval.

**Figure 1.**
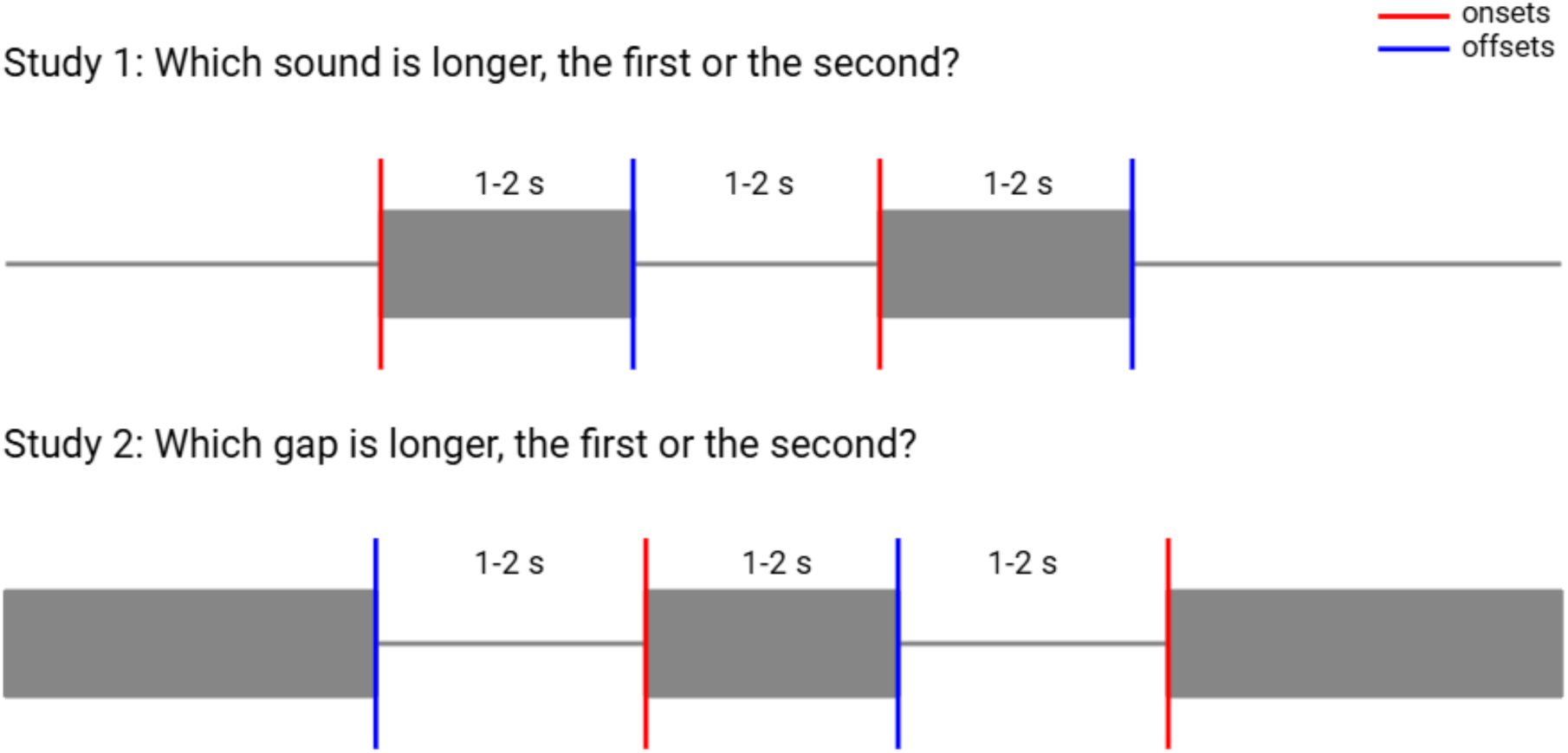
Experimental design for duration discrimination of noise intervals in silence (Study 1) and silent intervals (gaps) in noise (Study 2). Each study consisted of 2 sets of active and passive blocks, with no response required for the passive blocks. Schematics illustrate key features of the experimental design used to ensure offset responses were not obscured: (a) intervals were long enough to elicit offset responses; (b) the behavioural tasks required attention to both onsets and offsets; (c) durations were randomised over a large range; and (d) the order of onsets and offset cues was reversed between the two studies.

### Study 1: Duration discrimination of noise intervals in silence

Each trial consisted of a 1-second silence at the start of the trial, a first noise of randomised 1-2 second duration, a central silent gap of randomised duration (1-2 s), a second noise of randomised duration in the same range (1-2 s), and a following silent period throughout the response time (up to 3 s) and enough additional time to make each trial 15 seconds long. The long period of silence at the end of each trial was intended to reduce repetition suppression and enhancement that can affect individuals to differing degrees (Chandrasekaran et al., 2012). Importantly, there was no indication to the participant of the transition from the silent background at the end of one trial to the silent background at the beginning of the next.

### Study 2: Duration discrimination of silent intervals in noise

Each trial consisted of a 1-second noise at the start of the trial (plus 3 additional seconds for the first trial), a first silent gap of randomised 1-2 second duration, a central noise of randomised duration (1-2 s) and a second gap of randomised duration (1-2 s), followed by a period of noise which extended throughout the response time (up to 3 s) and enough additional time to make each trial 15 seconds long. Again, there was no indication to the participant of the transition from the noise background at the end of one trial to the noise background at the beginning of the next.

Noise stimuli were 60 dB SPL Gaussian white noise, with 5 ms cosine-squared ramps at rise and fall. The durations of the stimuli were at least 1 second long to allow complete recovery from stimulus onset and to enable offset ERPs to be cleanly distinguished from onset ERPs. Trial times were long (10 seconds) to minimise habituation-related decreases in N100 amplitude. Durations of pauses between the two sounds were randomised over the same 1-2 s range as durations of noises, so that offsets and onsets were preceded by equivalent durations of noise or silence.

For each participant, an experimental session involved four blocks of trials, consisting of two matched pairs of passive listening and active duration discrimination blocks. Each block was 7.5 minutes long and consisted of 30 trials, and the exact durations of randomised noise and silent intervals used for the trials were matched within pairs of passive and active blocks. The order of block presentation was alternated, so for the first pair of matched blocks, the passive block preceded the active block, and for the second pair, the passive block followed the active block. This design provided some counterbalancing of passive and active blocks; enabled the participant to become comfortable with the stimulus before responding in the first active block; and ensured that active blocks were performed in the middle of the session, rather than at the end when participants might be most tired.

The participant was seated in a soundproof booth in a comfortable chair with a keyboard support, facing a display screen. Short breaks (typically 5-10 minutes) were provided between blocks to alleviate fatigue. In passive listening blocks, the participant was instructed to focus on a cross at the centre of the screen in front of them while the sounds were presented. In active duration discrimination blocks, the participant was asked to fixate on the cross while the sounds were presented and then to respond at the end of each trial with a key press to indicate whether the first or the second sound was longer. Participants were instructed to respond in a few seconds following the sounds, but they were not required to respond quickly, as it was more important that they were accurate.

The auditory stimuli were presented diotically through EarTone in-ear earphones from a stimulus computer using the Psychophysics Toolbox (Brainard, 1997) extension in MATLAB (The MathWorks, Inc.). A 64-electrode Biosemi system (Biosemi Active Two AD-box ADC-17, Biosemi, Netherlands) was used to acquire the EEG data, with electrode placement following the 10-10 system. The amplifier sampling rate was 2048 Hz with a 0.1-100 Hz band pass filter. Voltage offsets from the reference electrode CMS were kept within +/- 20 mV at each electrode. The data were pre-processed and analysed using EEGLab and LIMO EEG toolboxes (Delorme and Makeig, 2004; Pernet et al., 2011) for MATLAB.

For analysis, the data were re-referenced to the two mastoid electrodes and downsampled to 256 Hz. Signals were high-pass (1 Hz, order 846) and low-pass (20 Hz, order 170) filtered using zero-phase Hamming-windowed sinc FIR filters (-6 dB cutoff frequency of 0.5 Hz for high-pass and 22.5 Hz for low-pass). Non-normally distributed data were detected based on kurtosis values (5 standard deviations from the average kurtosis), and any rejected electrode channels were interpolated. Independent component analysis (ICA) was used to manually identify and remove the component containing blink artefacts from the complete dataset.

Following these steps, sound onset and sound offset event epochs (from 0.5 s prior to 1 s after the event) were extracted for both passive and active blocks. This process yielded 60 onset-related and 60 offset-related epochs for each block, and a total of 120 epochs each for onset/offset events per active or passive block for each participant session. Baselines were removed for the epochs using the 0.5 s period before the event.

Event-related potentials (ERP) were investigated by averaging the epochs across blocks and participants (grand-average). A Bayesian bootstrap estimation of the 20% trimmed means with 95% HDI (highest density intervals) was used to generate ERP plots and difference plots between ERPs (Rousselet et al., 2023; Wilcox & Rousselet, 2023). The HDI is the confidence interval of the mean, and in the case that the HDI includes zero, i.e. overlaps with the x-axis, we can accept the null hypothesis instead of merely failing to reject it.

## Results

Study 1 (noise intervals in silence) and Study 2 (silent intervals in noise) both confirmed that offset ERPs are approximately one-third to one-half as large as onset ERPs for matched durations of noise and silent intervals. Table 1 shows the ERP component amplitudes and latencies for active duration discrimination and passive listening conditions in both studies. In the Study 1 active condition, the magnitude of the P200 component of the offset ERP was just under half that of the onset ERP (ratio of 0.45), with an amplitude difference of 5.71 µV. This ratio was reduced to 0.37 in the active condition of Study 2, with an amplitude difference of 5.97 µV. Another possible comparison is of the magnitude from N100 to P200. In the active condition of Study 1, the ratio of N100-P200 magnitudes between offset ERP (4.62 µV, from baseline as there was no N100) and onset ERP (12.79 µV) was 0.36. In Study 2, when the functional roles of onset and offset cues as interval start/end markers were reversed, the ratio of N100-P200 magnitudes between offset ERP (6.53 µV) and onset ERP (11.87 µV) was 0.55.

**Table 1.** P50, N100 and P200 magnitudes [95% HDI] and latencies for onset and offset ERPs in Study 1 (noise intervals in silence) and Study 2 (silent intervals in noise). The P200 component was reliably observed in offset as well as onset ERPs in both studies and across both active duration discrimination and passive listening conditions. Within each study and condition, this common P200 component was consistently one-third to one-half as large in the offset ERP as in the onset ERP. Latency of the P200 component was longer and amplitude slightly higher in active than passive conditions for ERPs corresponding to end-interval transients (offset ERP in Study 1, onset ERP in Study 2). There was no P50 in any condition for the offset ERP. There was no N100 (but a P100 instead, see Figure 4) for the offset ERP in the active condition of Study 1.

| *Study 1* | **P50** |  | **N100** |  | **P200** |  |
| --- | --- | --- | --- | --- | --- | --- |
| | Magnitude ($\mu\text{V}$) | Latency (ms) | Magnitude ($\mu\text{V}$) | Latency (ms) | Magnitude ($\mu\text{V}$) | Latency (ms) |
| Active Onset | 2.62 [1.98, 3.26] | 54.7 | -2.46 [-3.21, -1.73] | 105.5 | 10.33 [9.57, 11.03] | 195.3 |
| Passive Onset | 1.64 [1.07, 2.31] | 54.7 | -3.69 [-3.00, -2.46] | 105.5 | 10.2 [9.41, 10.95] | 195.3 |
| Active Offset | - | - | - | - | 4.62 [3.81, 5.35] | 207.0 |
| Passive Offset | - | - | -1.16 [-1.86, -0.53] | 66.4 | 2.92 [2.26, 3.63] | 191.4 |

**Table 1. P50, N100 and P200 magnitudes [95% HDI] and latencies for onset and offset ERPs in Study 1 (noise intervals in silence) and Study 2 (silent intervals in noise).**
| *Study 2* | **P50** |  | **N100** |  | **P200** |  |
| --- | --- | --- | --- | --- | --- | --- |
| | Magnitude ($\mu\text{V}$) | Latency (ms) | Magnitude ($\mu\text{V}$) | Latency (ms) | Magnitude ($\mu\text{V}$) | Latency (ms) |
| Active Onset | 0.84 [-0.15, 1.73] | 62.5 | -2.41 [-3.46, -1.53] | 105.5 | 9.46 [8.41, 10.43] | 183.6 |
| Passive Onset | 1.16 [0.11, 2.00] | 62.5 | -2.52 [-3.00, -2.46] | 105.5 | 9.16 [8.03, 10.04] | 179.7 |
| Active Offset | - | - | -3.03 [-3.92, -2.19] | 85.9 | 3.50 [2.50, 4.47] | 246.1 |
| Passive Offset | - | - | -3.31 [-4.17, -2.32] | 85.9 | 3.51 [2.42, 4.58] | 246.1 |

### Onsets and offsets evoke ERPs with distinct waveforms

To compare the morphology of onset and offset ERP waveforms, we focused on the grand-average ERPs for onsets in Study 1 and offsets in Study 2 in the active conditions. This approach ensured that the participants were attending to the transients and that cognitive factors related to transient cue order were equivalent. Figure 2 shows the onset ERP measured in Study 1 (noise intervals in silence, where onset is the start-interval cue) and the offset ERP measured in Study 2 (silent intervals in noise, where offset is the start-interval cue). All group grand-average ERP plots use robust estimators with 95% HDI based on Bayesian bootstrap estimates. The topographic plots show that the maximal N100 and P200 peaks occurred at the fronto-central peaks, so we used the FCz electrode, where the signal-to-noise ratio was also the highest, for the grand-average ERP plots.

**Figure 2.**
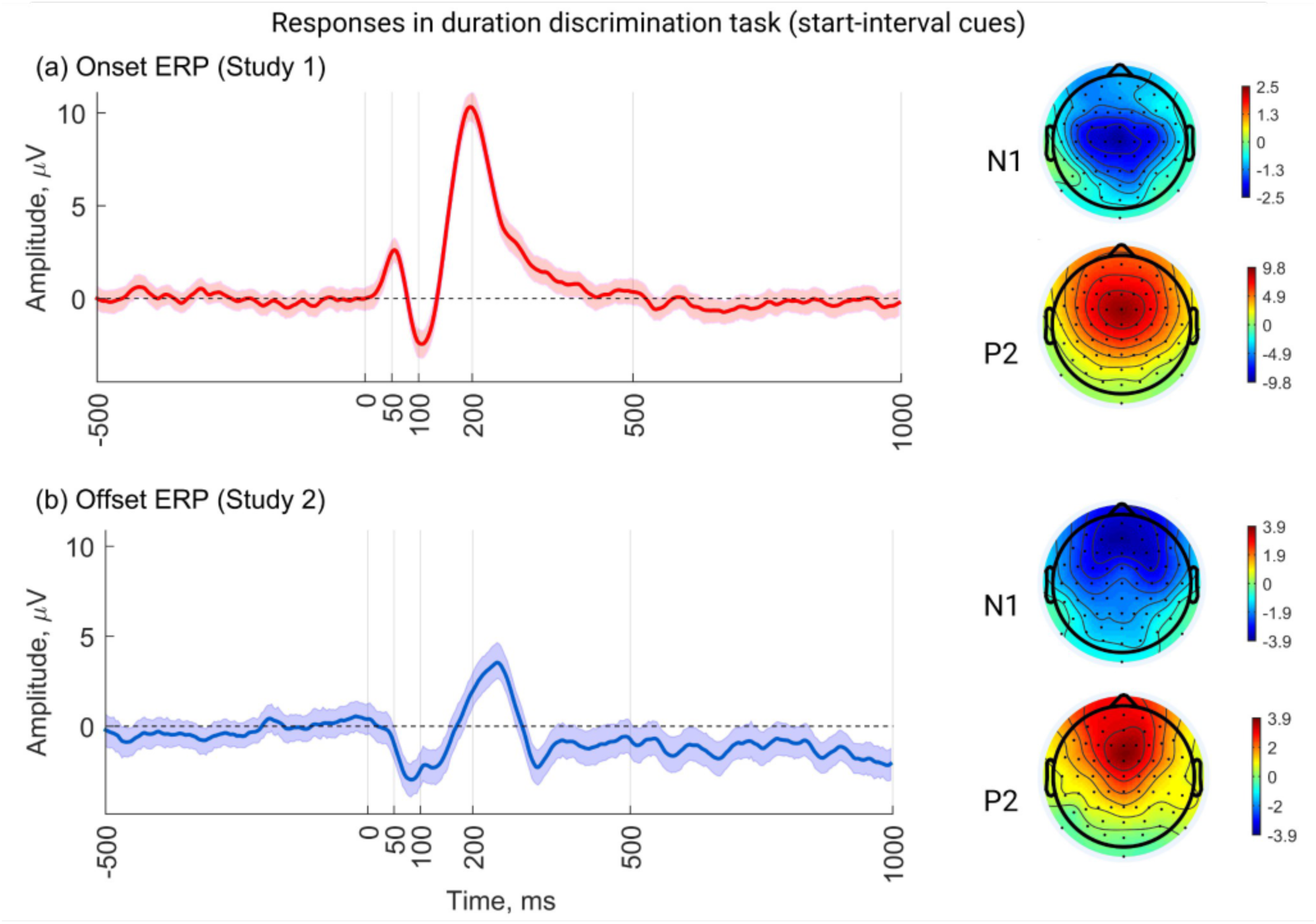
Sound onsets at the start of noise intervals (Study 1) and offsets at the start of silent intervals (Study 2) elicit ERPs with distinct waveforms. Plots show averages across trials in duration discrimination (active listening) blocks only. The onset waveform follows the typical ERP morphology, while there is no P50 component for the offset, and the fronto-central N1 occurs earlier at 85.9 ms with a second smaller peak at 121.1 ms. The offset P200 is delayed to 246.1 ms and has an amplitude of approximately a third of that of the onset P200. Topographic plots are shown for the maximal peaks. Error bars, 95% HDI.

The FCz grand-average ERP plots revealed several distinctive features of offset versus onset ERPs (Figure 2). First, there was no P50 component observed for the offset ERP, while the onset response followed the typical P50-N100-P200 waveform with P50 of 2.62 µV (95% HDI [1.98, 3.26]) at 54.7 ms and N100 of -2.46 ([-3.21, -1.73]) at 105.5 ms. Second, there was a double peak for the N100 in the offset response at 85.9 and 121.1 ms. The first peak in the complex (-3.03 µV [-3.92, -2.19]) was recorded as the N100, as it was the earliest, slightly larger magnitude, closest to 100ms, and also occurred at FCz, while the activity at the second peak (-2.38 µV [-3.29, -1.42]) was centred in the right temporal lobe electrodes. Third, there was a delayed P200 component for the offset ERP, at 246 ms, compared to the onset ERP at 195 ms, and the magnitude of the offset P200 (3.50 µV, 95% HDI [2.50, 4.47]) was approximately a third of the onset P200 magnitude (10.33µV [9.57, 11.03]). There was also a residual negativity in the offset ERP following the P200 activity that was absent for the onset response. Consistent with these observations, difference plot HDIs (not shown) revealed significant differences between the onset and offset ERPs as the initial cue of the durations in the P50 region (32 - 93 ms), P200 region (125 - 249 ms) and post-P200 activity (269 - 425 ms, 574 - 595 ms, 851 - 879 ms). There was no significant difference between onset and offset ERPs at the latency of the onset N100 (105.5 ms) based on the amplitude difference HDIs, despite the presence of a double-peaked rather than single-peaked N100 in the offset ERP.

### In duration discrimination tasks, sound offsets elicit either a P100 or N100 depending on behavioural context

Further investigation of onset and offset responses during active duration discrimination tasks in Study 1 and Study 2 revealed effects of behavioural context and cue order, particularly for offset ERPs (Figure 3). In Study 1 (when the offset cue was the end-interval event), the offset ERP exhibited a P100 component of 1.72 µV (95% HDI [0.99, 2.43]) at 105.5 ms. In Study 2 (when the offset cue was the start-interval event), the offset ERP instead exhibited a double-peaked N100 complex, with peaks of -3.03 µV [-3.92, -2.19] at 85.9 ms and -2.38 µV [-3.29, -1.42] at 121.1 ms. Thus, in duration discrimination tasks, both the polarity and the morphology of the P100/N100 wave in the offset ERP depended on whether or not the offset cue marked the start or the end of the discriminated interval.

**Figure 3.**
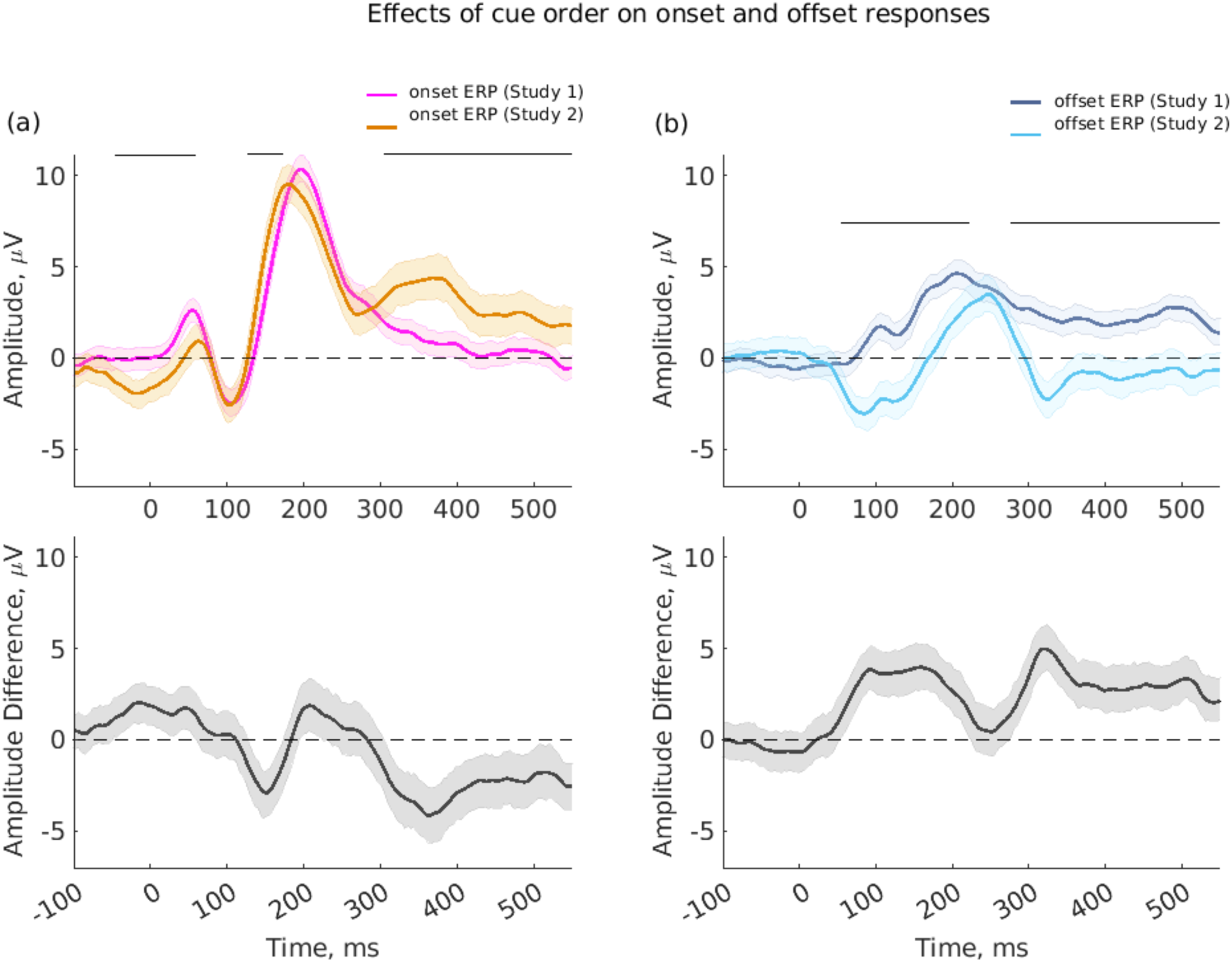
Dependence of onset and offset ERPs on behavioural context and cue order in active duration discrimination tasks. Top plots show onset ERPs (left) and offset ERPs (right) for Study 1 (noise intervals in silence) compared to Study 2 (silent intervals in noise). Bottom plots show ERP amplitude differences (Study 1 minus Study 2 ERPs). Post-P200 activity was enhanced for offsets in Study 1 and onsets in Study 2, indicating that the morphology of both onset and offset ERP waveforms depend in part on whether or not the transient serves as an end-interval event marker in the duration discrimination task. Offset ERPs additionally exhibited a reversal of polarity of the ERP around 100 ms depending on the nature of the duration discrimination task. In Study 1, when offsets were end-interval events, a P100 was observed in the offset ERP; in Study 2, when offsets were start-interval events, the offset ERP had a double-peaked N100 complex. Error bars indicate 95% HDI; horizontal floating lines in upper plots illustrate where amplitude differences are significantly different from zero.

Additionally, when offsets were the end-interval event (i.e., end of noise intervals in Study 1), post-P200 activity was observed in the offset ERP, with a delayed P400 peak of 2.78 µV [2.08, 3.39] at 484.4 ms. Similar post-P200 activity was observed in the onset ERP when onsets were the end-interval event (i.e., end of silent intervals in Study 2), with a P400 peak of 4.45 µV (95% HDI [3.28, 5.67]) at 371.1 ms.

However, unlike the offset responses, onset responses in both Study 1 (noise intervals in silence) and Study 2 (silent intervals in noise) maintained the characteristic P50-N100-P200 components for sensory detection, with no reversals of polarity. The ERP difference waveforms (Figure 3, bottom row) confirmed significant differences between Study 1 and Study 2 for post-P200 activity in both onset and offset ERPs, but significant differences at the ∼100 ms latencies of N100/P100 only for offset ERPs.

Latencies of ERP components also revealed effects of behavioural context in duration discrimination tasks. Both onset and offset ERPs elicited faster brain responses when they were start-interval rather than end-interval cues. For example, in Study 1, when onsets were the start-interval cue, the latency of the P50 response in the onset ERP was 54.7 ms, compared to 62.5 ms in Study 2 (when onsets were the end-interval cue). For offsets, which generated no P50 response in either study, the earliest ERP peak was the N100/P100 component. In Study 2, offsets served as start-interval cues and generated an N100 complex with a double peak starting at 85.9 ms, while in Study 1, offsets served as end-interval cues and generated a P100 at 105.5 ms. Similarly, the latencies of P200 components in both onset and offset ERPs were shorter for start-interval than end-interval cues.

### Active listening modulates both early and late components of ERP waveforms for end-interval offsets but only late components for end-interval onsets

We further explored the impact of task engagement by comparing ERPs between active duration discrimination blocks and passive listening blocks within each study. Figure 4 shows the results for Study 1 (noise intervals in silence), where onsets were the start-interval cue and offsets the end-interval cue. For the offset ERP in the active condition, there was an additional P100 peak at 105.5 ms instead of an N100 at 66.4 ms in the passive condition, and the P200 component had a considerably larger peak (4.62 µV [3.81, 5.35] at 207.0 ms, compared to a slightly earlier P200 of 2.92µV [2.26, 3.63] at 191.4 ms in the passive condition). There was also greater post-P200 activity, presumably related to the duration judgement following the end-interval event. In contrast, for onset ERPs, differences in amplitude between active and passive conditions were small (maximum difference is 1.47 µV [0.44, 2.47] at 89.8 ms) and the waveform morphology remained relatively stable. These observations are further illustrated in the difference plots, which reveal significant amplitude differences between active and passive conditions primarily for offset ERPs, with difference waveform peaks at latencies of 101.6 ms (1.79 µV [0.82, 2.75]) and 269.5 ms (2.80 µV [1.89, 3.88]).

**Figure 4.**
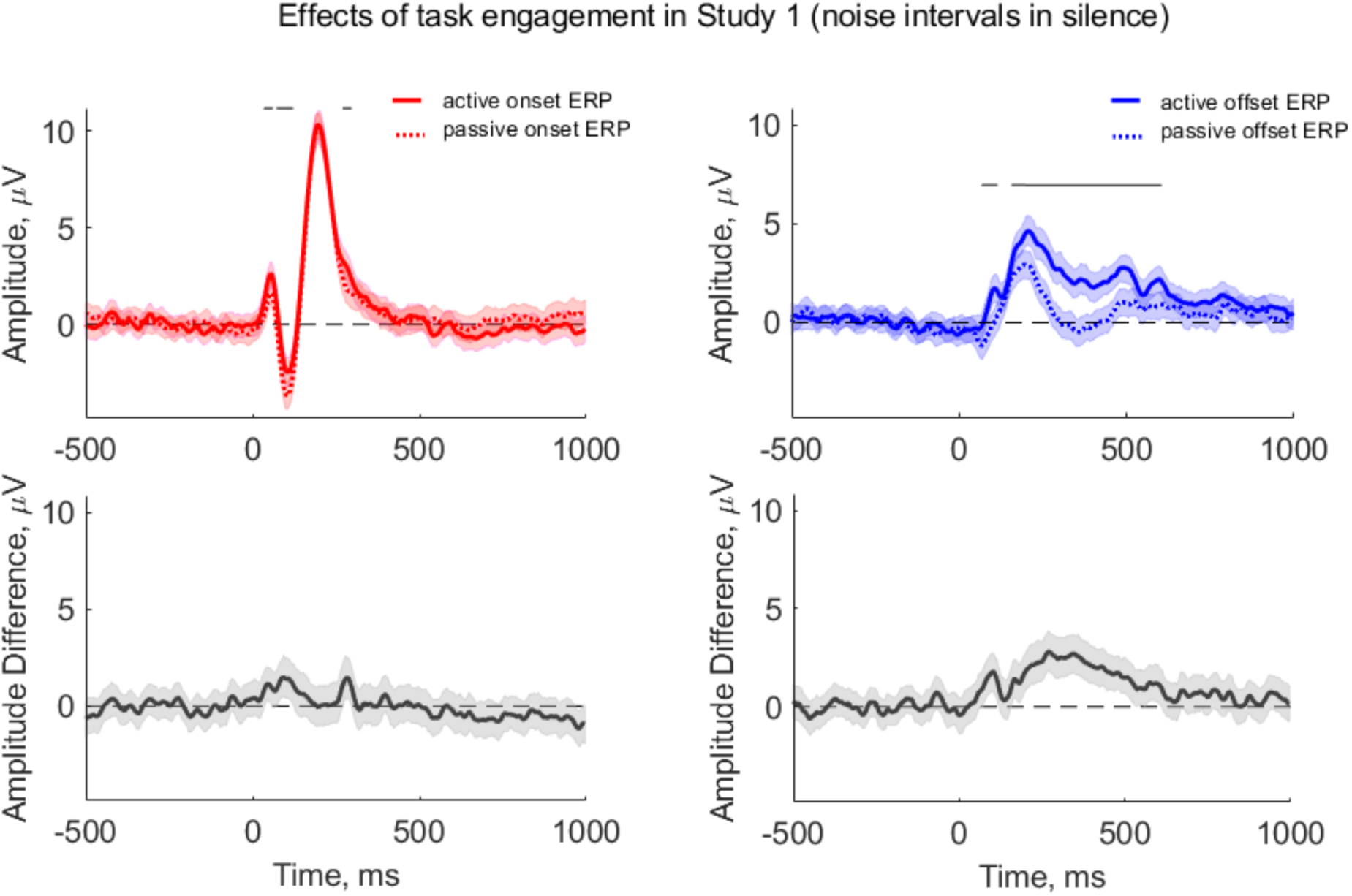
Differences between active duration discrimination and passive listening conditions appear in both early and late components of offset ERP waveform in Study 1 (noise intervals in silence). Top plots show onset ERPs (left) and offset ERPs (right) for active duration discrimination and passive listening conditions (solid and dotted lines, respectively). Bottom plots show ERP amplitude differences (active minus passive condition ERPs). For offsets, which were the end-interval event in Study 1, there were significant differences in ERP morphology between active and passive conditions: a P100 peak in the active condition instead of the small N100 in the passive condition; differences in the magnitude of the P200 peak; and prolonged differences in post-P200 activity. For onsets, differences in ERP amplitude between the active and passive conditions were much smaller and less extended in time (at 36-58 ms, 71-124 ms and 271-297 ms). Conventions for error bars and significance lines as in Figure 3.

Figure 5 shows results for Study 2 (silent intervals in noise), when the functional roles of onsets and offsets were reversed; offsets were the start-interval cue and onsets the end-interval cue. In this case, prominent amplitude differences between active and passive blocks were observed for the onset ERP rather than the offset ERP, but only in the post-P200 period (240-572 ms after the end-interval onset). The start-interval offset ERPs exhibited relatively similar morphology and amplitude in active and passive blocks, with only a small difference (peak of 2.61 µV [1.30, 4.06] at 351.6 ms) in the regions 332-380 ms and 438-461 ms owing to a greater undershoot in the passive condition.

**Figure 5.**
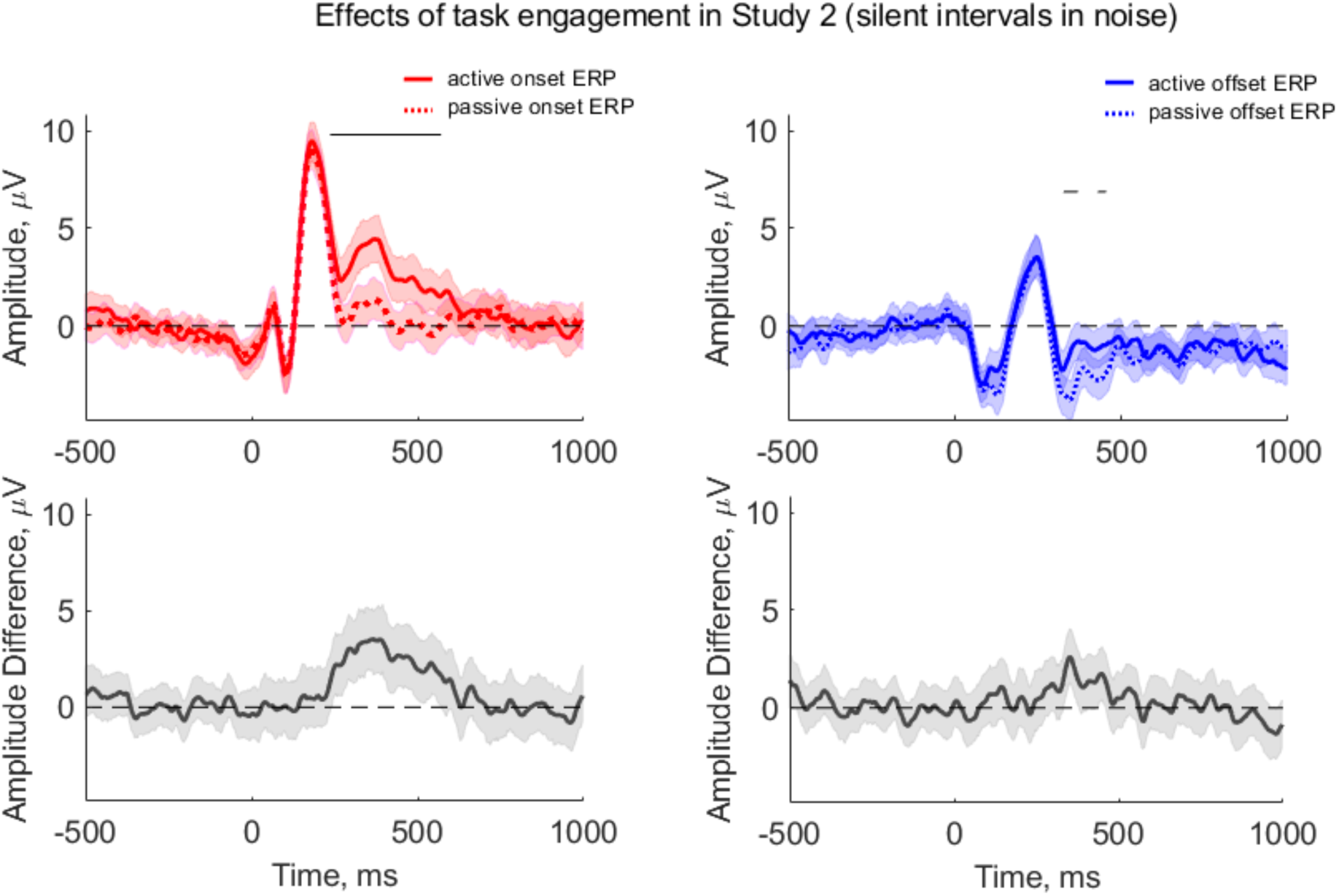
Differences between active duration discrimination and passive listening conditions appear primarily in late components of the onset ERP waveform in Study 2 (silent intervals in noise). All plot conventions as in Figure 4. For onsets, which were the end-interval event in Study 2, there were significant differences in ERP amplitude between 240 and 572 ms following the transient. For offsets, there were small differences at 332-380 ms latency owing to greater negativity post-P200 in the passive condition.

In summary, late ERP components (after P200) were enhanced in active compared to passive blocks for offsets in Study 1 and for onsets in Study 2, presumably due to cognitive processes related to judgement of duration after the end-interval event in active blocks. For offsets, however, ERP differences between active and passive blocks were also evident in the pre-P200 period, especially around the typical N100, where a P100 component present in the active condition was absent in the passive condition. No differences between active and passive conditions were observed in such an early time window for the onset responses. These results indicate that engagement in a duration discrimination task specifically alters the P100/N100 response to offsets when they are end-interval events, and more generally enhances post-P200 activity for any end-interval event (onset or offset).

## Discussion

Sound offsets are less salient than sound onsets, and the physiological mechanisms that generate responses to sound offsets in the brain are thought to be at least partly dissociable from the mechanisms generating responses to sound onsets (Scholl et al., 2010; Anderson & Linden 2016; Kopp-Scheinpflug et al. 2018; Anandakumar and Liu 2022; Edeline & Liu 2025). Previous EEG studies of onset and offset responses in the human brain have identified asymmetries in onset and offset ERPs (Onishi & Davis, 1968; Spychala et al., 1969; Hillyard et al. 1978; Pantev et al., 1996; Pratt et al., 2008; Volosin & Horvath, 2020), as expected given the perceptual and mechanistic asymmetries. However, potential confounds in previous studies, such as temporal predictability of onsets and offsets, have left many unresolved questions about the true nature of asymmetries in onset and offset ERPs.

Results presented here identify distinctive and reliable features of the sound-offset-related ERP profile in humans and resolve open questions about asymmetries in morphology and task-dependence between onset and offset ERPs. Several features of the experimental design helped to overcome potential confounds in previous work. By requiring participants to discriminate the duration of either noise intervals in silence or silent intervals (gaps) in noise, we were able to explore the role of behavioural context and cue order in shaping onset and offset ERPs. By presenting exactly the same stimuli in active duration discrimination and passive listening conditions for each participant, we could analyse effects of task engagement on onset and offset ERPs. By using noise and silent intervals of 1-2 s duration in all experiments, we could cleanly distinguish onset and offset ERPs with no overlap. Finally, by randomising both noise and silent interval durations across many finely discretized possibilities within the 1-2 s range, we minimised and equalised the temporal predictability of onset and offset events and ensured that the duration discrimination task could not be performed without close attention to transient timing.

Our results show that even when the functional roles of onsets and offsets were reversed (i.e., offset cues were at the start rather than end of a discriminated interval), offset ERPs were significantly smaller than onset ERPs. The most prominent component in all conditions tested was the P200, and this had a similar topography for both onset and offset responses, with maximal amplitudes at the fronto-central electrodes. The amplitude of the P200 was approximately one-third to one-half as large in offset ERPs as onset ERPs within and across participants, conditions and studies. These amplitude differences are consistent with those observed in previous studies using tones rather than noise stimuli (Onishi and Davis, 1968; Spychala et al., 1969), and may therefore be robust features of offset responses across stimulus and behavioural conditions, at least when preceding periods of sound or silence are equivalent (cf. Pfefferbaum et al., 1971).

Offset ERPs were also morphologically different from onset ERPs. Offset responses did not follow the characteristic P50-N100-P200 waveform typical of onset responses to sensory stimuli. There was no P50 for the offset ERP response in any condition, consistent with previous reports (Hari et al., 1987; Pantev et al., 1996; Yamashiro et al., 2009; Pratt et al., 2008; Zhang et al., 2016). Moreover, the offset response around 100 ms latency was morphologically complex and dependent on the behavioural context in which the offset occurred. When the offset was the end-interval event in an active duration discrimination task, a P100 was observed. However, when offsets were the start-interval events in a duration discrimination task, or either the start-interval or end-interval event in a passive listening task, the offset ERP measured at FCz had a double-peaked N100 wave. In contrast, in onset ERPs, the basic morphology of the P50 and N100 waves remained relatively stable across stimulus and behavioural conditions. Thus, early waves of the offset ERP appear to be more dependent on stimulus and behavioural context than early waves of the onset ERP. This asymmetry likely reflects differences in the generators of early waves of onset and offset ERPs. Indeed, previous studies of the double-peaked N100 in offset ERPs found that source localisation of the earlier component was more frontal and resembled that of N100 in onset ERPs, while source localisation for the later component was more central/temporal (Pratt et al., 2005; Pratt et al., 2007).

Interestingly, like the P100 response in the offset ERP, post-P200 activity in both onset and offset ERPs was dependent on engagement in a duration discrimination task and on whether the transient marked the start or the end of the interval for which duration was being judged. Post-P200 activity for both onsets and offsets exhibited larger amplitude and longer latency when the transient was the end-interval event in an active duration discrimination task, compared to when it was the start-interval event in a duration discrimination task or either the start- or end-interval event in a passive listening task. These observations suggest that post-P200 activity is related to decision-making in the duration discrimination task and is not specific to one type of transient event. The findings for post-P200 activity also serve to highlight the asymmetries between offset and onset ERPs in the much earlier P100/N100 wave, where a similar apparent sensitivity to decision-making in the duration discrimination task was evident only in the offset ERP, not in the onset ERP.

The involvement of early ERP components in the duration discrimination judgement for offset but not onset end-interval cues suggests that top-down mechanisms play a greater role in offset than onset perception. Precise offset detection may require more attention than onset detection, given that sound onsets generally elicit stronger neurophysiological responses regardless of task demands and active or passive listening. This observation has ecological validity, as onset responses are alerting to potential dangers in the environment and often indicate the appearance of a new object in the auditory scene (Bregman, 1990). In the first EEG study on auditory responses, Davis described how increased drowsiness enhanced the onset response (Davis, 1939). The automacy of onset responses is in contrast to the offset response, which has been reported to be much more salient in awake animals than when barbiturate anaesthetics are used (Qin et al, 2007; Zurita et al., 1994). Thus, although both onsets and offsets are equivalently low-level sound features, onset detection likely involves preattentive processing including arousal responses, while active listening may be required for a sound offset to cause arousal.

Overall, this work adds to growing evidence for fundamental differences in the neurophysiological mechanisms of onset and offset detection (Scholl et al., 2010; Anderson & Linden, 2016; Kopp-Scheinpflug et al., 2018; Anandakumar & Liu, 2022; Edeline & Liu, 2025) and establishes a simple and robust experimental paradigm for further development of offset ERP measurement as a potential clinical tool (cf. Colak et al., 2025). A previous study has shown that at least in mice, auditory temporal processing abnormalities can arise from specific deficits in sound-offset processing that do not alter central auditory responses to sound onsets (Anderson & Linden, 2016). Poor brain sensitivity to sound offsets has also recently been shown to correlate with speech-in-noise perception difficulties in older adults (cf. Colak et al., 2025). Poor offset sensitivity, arising either from central auditory abnormalities in offset processing pathways or more cognitive factors such as impaired ability to attend effectively to temporal cues, would not be detected on standard audiometric tests but could contribute to auditory temporal processing disorders including difficulties with gap detection, duration discrimination and speech perception (Füllgrabe et al., 2015; Kopp-Scheinpflug and Tempel, 2015; Kopp-Scheinpflug et al., 2018). Better assessment of offset sensitivity in humans using careful experimental design and temporally precise tools such as EEG may lead to improved diagnostics and treatment for difficulties with auditory temporal processing and speech perception that are currently poorly understood.

## Additional Information

### Data availability

Data are available from the corresponding author upon request.

### Author contributions

FA: Conceptualization, Investigation, Data Curation, Software, Visualization, Formal Analysis, Writing - Original Draft, Funding Acquisition.

GB: Investigation, Data Curation, Software, Formal Analysis.

AS: Visualization, Formal Analysis, Writing - Review & Editing.

JL: Conceptualization, Supervision, Visualization, Formal Analysis, Writing – Review & Editing, Funding Acquisition.

## Acknowledgements and funding.

This work was supported by a PhD studentship from the UCL Engineering and Physical Sciences Research Council Doctoral Training Programme (FA); by a research grant from the Medical Research Council and Wellcome Trust via the UCL Therapeutic Acceleration Support Fund (JL); and by research funding from the NIHR-UCLH Biomedical Research Centre Deafness and Hearing Problems / Hearing Health Theme (JL).

